# Quantitative mapping of synaptic periactive zone architecture and organization

**DOI:** 10.1101/2022.06.16.496425

**Authors:** Steven J. Del Signore, Margalit G. Mitzner, Anne M. Silveira, Thomas G. Fai, Avital A. Rodal

## Abstract

Following exocytosis at active zones, synaptic vesicle membranes and membrane-bound proteins must be recycled. The endocytic machinery that drives this recycling accumulates in the periactive zone (PAZ), a region of the synapse adjacent to active zones, but the organization of this machinery within the PAZ, and how PAZ composition relates to active zone release properties remains unknown. The PAZ is also enriched for cell adhesion proteins, but their function at these sites is poorly understood. Here, using Airyscan and STED imaging of *Drosophila* synapses, we develop a quantitative framework describing the organization and ultrastructure of the PAZ. Different endocytic proteins localize to distinct regions of the PAZ, suggesting that sub-domains are specialized for distinct biochemical activities, stages of membrane remodeling, or synaptic functions. We find that the accumulation and distribution of endocytic but not adhesion PAZ proteins correlate with the abundance of the scaffolding protein Bruchpilot at active zones - a structural correlate of release probability. These data suggest that endocytic and exocytic activities are spatially correlated. Taken together, our results provide a new approach to quantify synaptic architecture and identify novel relationships between the exocytic and endocytic apparatus at the synapse.

## 2. Introduction

Endocytic proteins fulfill multiple physiological functions at the synapse, including synaptic vesicle endocytosis and recycling, transmembrane protein trafficking, release site clearance, and cargo sequestration (Maritzen and Haucke, 2018). The dozens of endocytic proteins that drive these diverse functions accumulate at micromolar concentrations at synapses (Wilhelm et al., 2014) and in many cases are pre-deployed to the plasma membrane (Del Signore et al., 2021; Imoto et al., 2021) in a poorly defined micron-scale domain called the periactive zone (PAZ) (Cano and Tabares, 2016; Roos and Kelly, 1999; Sone et al., 2000). This is in sharp contrast to non-neuronal cells, where the same endocytic proteins are recruited to diffraction limited membrane spots in a well-defined temporal sequence (Kaksonen and Roux, 2018). These unique features of endocytic proteins at synapses highlight the need to investigate the synapse-specific mechanisms that control endocytic protein recruitment and activity. However, the imaging-based approaches that have been so useful in other cell types for measuring actions of sequentially recruited endocytic proteins are incompatible with the high concentration and constitutive pre-deployment of endocytic proteins at the synapse. Thus, it remains difficult to investigate how synapses control where and when endocytic molecules are active relative to sites of exocytosis, and how the recruitment and activation of dozens of endocytic proteins are tailored to their many synaptic functions, particularly the trafficking of diverse cargoes.

Despite the high concentration of endocytic proteins at the PAZ, this region is not a monofunctional endocytic zone. Endocytosis can proceed through multiple distinct molecular mechanisms (e.g. Clathrin mediated, bulk, ultrafast, kiss-and-run) (Gan and Watanabe, 2018) and occur both within the PAZ and at/near AZs (Kidokoro, 2006; Koenig and Ikeda, 1996; Kosaka and Ikeda, 1983; Kuromi et al., 2010; Teng and Wilkinson, 2000; Watanabe et al., 2013a). Further, PAZ molecules have many functions beyond synaptic vesicle endocytosis, as mentioned above. Finally, in addition to endocytic proteins the PAZ contains a high concentration of synaptic adhesion molecules. Synaptic adhesion proteins regulate synapse formation, maintenance, and plasticity, and are implicated in multiple neurological disorders (Jang et al., 2017; Südhof, 2018; Sytnyk et al., 2017). They are especially interesting to consider as PAZ constituents because they regulate endocytosis and synaptic vesicle trafficking (Dagar et al., 2021; Leshchyns’ka et al., 2006; Luo et al., 2021; Shetty et al., 2013; van Stegen et al., 2017), and are themselves endocytic cargoes. Indeed, presynaptic endocytosis and trafficking pathways regulate the levels and function of adhesion molecules (Bailey et al., 1992; Blanchette et al., 2022; Fu and Huang, 2010; Kurshan et al., 2018; Nahm et al., 2016; Yamada et al., 2007). Therefore, in addition to being important components of the PAZ, adhesion molecules are crucial regulators and effectors of the endocytic machinery at the synapse. Our goal in the current work is to develop an approach to quantify the distribution of endocytic and adhesion molecules at the PAZ to begin to provide insight into their different functions.

The elaborate organization of endocytic and adhesion molecules into a PAZ domain is conserved in many synapse types across species. It has been most frequently described at large synapses such as the *Drosophila* neuromuscular junction (NMJ; Wan et al., 2000; Sone et al., 2000; Roos and Kelly, 1999; Estes et al., 1996; Schuster et al., 1996; Koh et al., 2004; González-Gaitán and Jäckle, 1997; Marie et al., 2004; Rodal et al., 2008), photoreceptor ribbon synapses (Gray and Pease, 1971; Wahl et al., 2016, 2013), and lamprey reticulospinal axons (Bloom et al., 2003; Evergren et al., 2004). Endocytic proteins are also pre-deployed adjacent to active zones at small central synapses such as in hippocampal neurons (Gerth et al., 2017; Imoto et al., 2021). The evidence that the PAZ is a bona fide membrane domain derives from direct observations of endocytic membrane intermediates by electron microscopy (Heuser and Reese, 1973; Watanabe et al., 2013b), by the enrichment of endocytic or synaptic vesicle associated F-actin by EM and fluorescence microscopy (Bloom et al., 2003; Del Signore et al., 2021; Shupliakov et al., 2002), and by the accumulation of known endocytic proteins by fluorescence microscopy (Del Signore et al., 2021; Gerth et al., 2017; Imoto et al., 2021; Marie et al., 2004; Sone et al., 2000). While most studies have treated the PAZ as a homogenous and ill-defined structure, there are hints that some PAZ components, such as the adhesion protein FasciclinII (FasII), might accumulate in distinct subdomains around the active zone (Marie et al., 2004).

A further open question is how the spatial organization of the PAZ or other endocytic zones might be coupled to the exocytic machinery. At synapses generally, release sites are associated with active zones, which assemble to recruit and organize release machinery, synaptic vesicles, and voltage gated calcium channels (Ghelani and Sigrist, 2018; Petzoldt et al., 2016). As such, active zone composition and architecture are important determinants of synaptic physiological and homeostatic properties (Böhme et al., 2019; Fouquet et al., 2009; Goel et al., 2019; Hong et al., 2020; Kittel et al., 2006). For example, at the *Drosophila* NMJ, different active zones even within a single neuron exhibit up to 50-fold differences in release probability for both evoked and spontaneous release. These differences in release probability correlate with the age of the active zone, and with the composition and organization of active zone components including Brp/CAST and voltage-gated calcium channels (Akbergenova et al., 2018; Gratz et al., 2019; Melom et al., 2013). However, it remains unknown whether or how active zone assembly and physiology might control PAZ properties. This includes whether and how the endocytic machinery organizes developmentally (over minutes to hours) with respect to the release probability and composition of nearby active zones and how this organization responds acutely (in seconds) to exocytosis.

Despite all these data, no study to date has systematically and quantitatively analyzed the structure and molecular patterning of a PAZ or compared the properties of a PAZ with its associated exocytic machinery. This has been a challenge since the PAZ is heterogenous in shape and size (i.e. it is not a simple isotropic bullseye around the active zone), and its dimensions approach the limits of conventional light microscopy. Here we develop a new method to analyze the complex protein architecture of the synaptic endocytosis machinery. We apply this method to analyze the distribution of a subset of endocytic and adhesion proteins at the *Drosophila* NMJ (see Table 1), including Clathrin, Dap160, Nervous Wreck, Dynamin, and FasciclinII. We chose this panel for multiple reasons: (1) These proteins are well-studied molecules known to enrich at the PAZ at this synapse (2) Based on loss of function phenotypes, these molecules contribute to diverse synaptic functions including endocytosis, vesicle trafficking, and adhesion (3) The endocytic molecules selected are thought to act at distinct steps during endocytosis. This synapse is well-studied, has multiple active zones per synapse, well characterized physiological properties, and is highly amenable to super-resolution imaging. Further, the wide range of release probabilities between active zones at this synapse provide a large dynamic range to compare active zone and PAZ assemblies. With this approach, we identify several new features of PAZ architecture. First, we find that distribution of different PAZ proteins is highly heterogeneous, both within and between individual PAZ units. Second, we find that the overall structure of the PAZ is only partially defined by active zone position and pattern. Third, we find that the accumulation and distribution of specific PAZ proteins significantly correlate with the concentration of the scaffold BRP (a structural correlate of release probability) in nearby active zones. Together these findings suggest new relationships between active zone and PAZ molecular organization and pave the way for new studies investigating the molecular dynamics and mechanisms that couple the assembly and function of exocytic and endocytic assemblies at synapses.

**Table 1.**
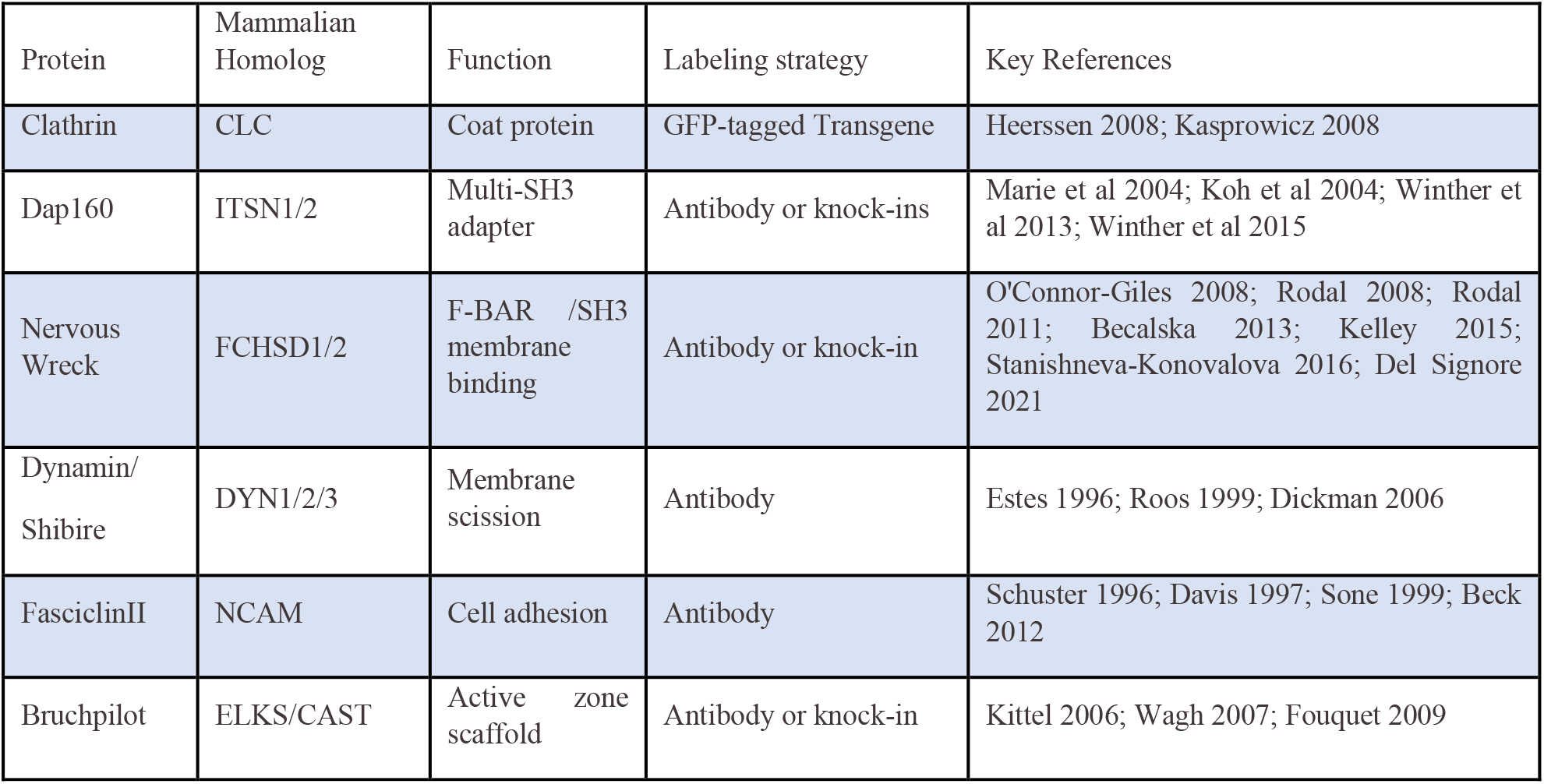
Summary of genes analyzed.

## 3. Materials and Methods

### *DROSOPHILA* IMAGING

#### *Drosophila* culture and strains

Flies were cultured using standard media and techniques. All flies were raised at 25°C. See Table 2 for fly strains used in this work.

**Table 2.**
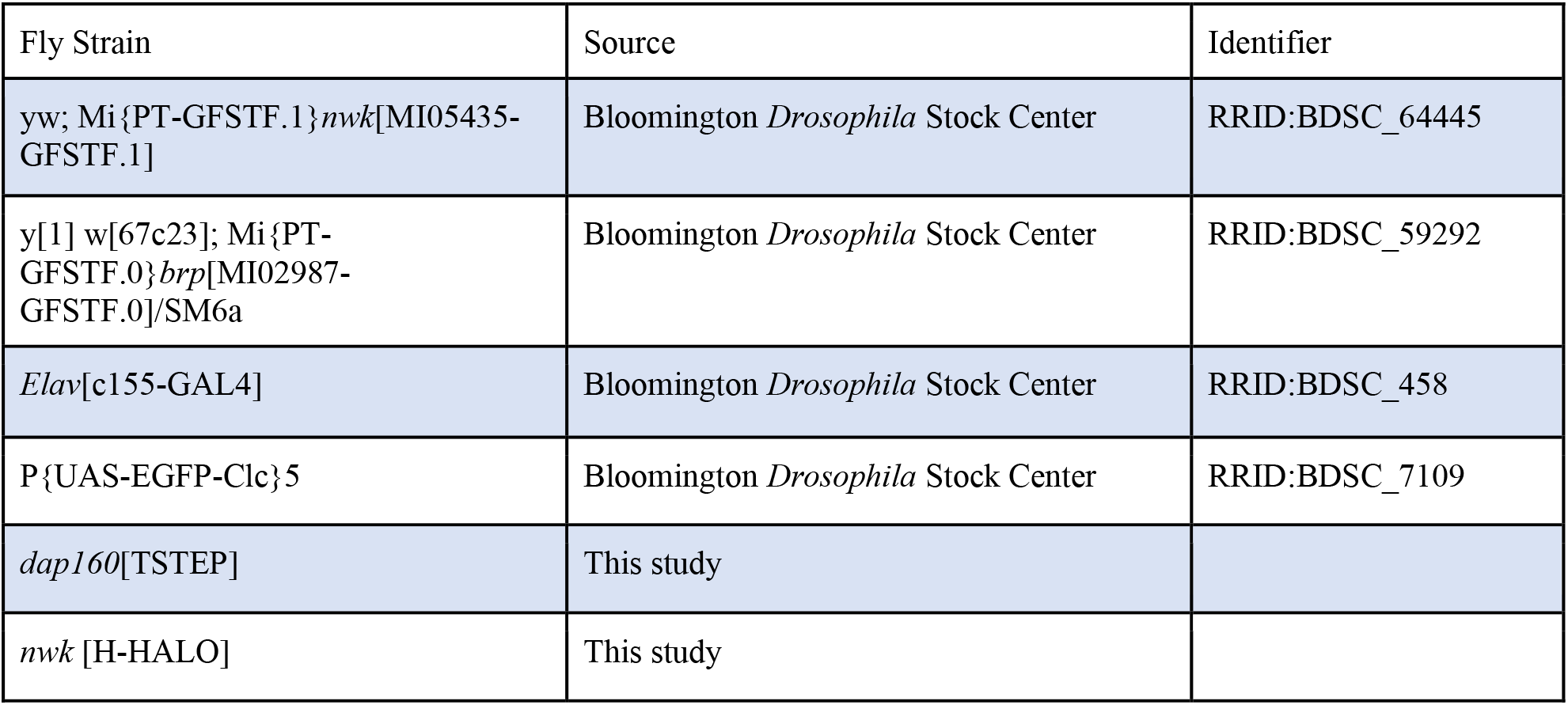
List of *Drosophila* strains used in this study.

#### Immunohistochemistry

For analysis of NMJ morphology and protein localization, flies were cultured at low density at 25°C. Wandering third instar larvae were dissected in calcium-free HL3.1 saline (Feng et al., 2004) and fixed for 20 min in HL3.1 containing 4% formaldehyde. Fixed larvae were blocked in 3% BSA in 0.1% Triton ×100 in PBS for 30-60 min and incubated with primary antibody in block solution either overnight at 4° C or 2 hours at room temperature, followed by washing and incubation with dye-conjugated secondary antibodies (Jackson Immunoresearch). Larvae were mounted in Prolong Diamond mounting medium. Primary antibodies used were rabbit anti-Nwk 970 (RRID:AB_2567353 (Coyle et al., 2004)), rabbit anti-Dap160 (RRID:AB_2569367 (Roos and Kelly, 1998)), rabbit anti-Dyn (RRID:AB_2314348 (Estes et al., 1996)), mouse anti-Dyn ((RRID:AB_397640, BD Biosciences Clone 41)), mouse anti-BRP (RRID:AB_2314866, DSHB clone nc82), mouse anti-FasII (RRID:AB_528235 (DSHB clone 1D4)). For Airyscan imaging, larvae were minimally stretched and mounted in an excess volume of mounting medium to preserve 3D morphology of boutons. For STED imaging, larvae were stretched and mounted in a minimum volume of medium to render preparations flat to aid 2D imaging. Live HALO labeling of Nwk::HALO was performed as follows: we dissected heterozygous Nwk::HALO/+ larvae in HL3.1 medium, incubated fillets in 250 nM HALO-JF549 (Promega, Inc.) for 5 minutes, washed in HL3.1 for five minutes, then mounted and imaged fillets immediately. All images were acquired within 15 minutes of mounting.

#### Image acquisition

Confocal and Airyscan images of NMJs were acquired with a Zeiss 880FAS microscope with a 63X (NA1.4) oil immersion objective in LSM or SR mode, respectively, using Zen Blue software. All raw image stacks were processed in Zen Blue to construct Airyscan images using 3D Airyscan processing with automatic settings. We quantified optical resolution using tetraspeck beads (Invitrogen) at 560nm as 158 × 179 × 401 nm in XYZ, respectively. We found this resolution to be sufficient to subdivide the PAZ, localize AZ and PAZ components, and generate radial profiles of PAZ units, and provide results consistent with STED microscopy (see **Fig 5**).

2D-STED images were acquired with an Abberior FACILITY line STED microscope with 60x (NA1.3) silicone immersion objective, pulsed excitation lasers (561 nm and 640 nm) and a pulsed depletion laser (775 nm) used to deplete all signals. BRP was labeled with STAR RED (Abberior, Inc.) and both Nwk and Dyn were labeled with STAR ORANGE (Abberior, Inc.). Pixel size was set to 30 nm, and single 2D slices were acquired of terminal branches of NMJs that lay in a single focal plane. All images within one dataset were acquired with the same microscope settings (see Supplemental Table 1 for N).

#### Generation of Dap160^KI^

The endogenous dap160 locus was tagged with a tissue-specific convertible TagRFPt to EGFP tag (T-STEP (Koles et al., 2015)). Briefly, the following genomic sequence was used to target DNA cleavage by Cas9: ATGTGAGATTCACTTCTTGGTGG (at location 2L:21139297--21139319). The T-STEP cassette was flanked with 5’ homology (2L:21138474-21139307) and 3’ homology (2L:21139308-21140194) arms, and gene targeting was carried out as described ((Koles et al., 2015), based on (Chen et al., 2015)). Dap160-TSTEP flies were crossed to C380-Gal4;;UAS-Rippase::PEST @attP2 flies to isolate motor neuron ripouts of the TagRFPt cassette, leaving a C-terminal EGFP knockin.

#### Generation of Nwk-Halo

A HALO7 tag was knocked into the endogenous locus by CRISPR-CAS9 mediated gene editing (performed by Wellgenetics, using modified methods of (Kondo and Ueda, 2013)). In brief, gRNA sequence AGCCACGGACAGTTATAGAG[CGG] was cloned into U6 promoter plasmid(s). This targets the stop codon in exon 22, thus labeling isoforms H,I,J, and L. Cassette Halo7-3xP3-RFP, which contains Halo7 and a floxed 3xP3-RFP, and two homology arms were cloned into pUC57-Kan as donor template for repair. nwk/CG43479-targeting gRNAs and hs-Cas9 were supplied in DNA plasmids, together with donor plasmid for microinjection into embryos of control strain w[1118]. F1 flies carrying selection marker of 3xP3-RFP were further validated by genomic PCR and sequencing. CRISPR generates a break in nwk/CG43479 and is replaced by cassette Halo7-3xP3-RFP. Finally, the floxed 3xP3-RFP cassette was excised to generate the knockin strain used in experimental crosses.

### PAZ PROCESSING AND SEGMENTATION

All image processing and data extraction were performed using custom FIJI and Python scripts (available: https://github.com/rodallab/paz-analysis).

#### Image pre-processing

Prior to PAZ segmentation and analysis, 2D maximum intensity projections of the bottom half of NMJ boutons were made semi-manually using a custom FIJI script as follows: ROIs were drawn manually around boutons lying in the same approximate Z plane, and the medial Z-slice was set. Maximum intensity projections of the lower portions of the stack were generated and saved, and the medial slice was also saved. As needed, images were manually ‘cleaned’ to remove axon or other spurious signal not belonging to a Type 1b motor neuron.

#### NMJ Segmentation

Segmentations of the entire bouton region(s) within each image were performed as follows: 1. Whole stacks were background subtracted (rolling ball radius = 50 pixels). 2. Channel intensities were normalized by the whole-image average value per channel. 3. All channels were summed to create a composite NMJ image. The composite image was blurred using a Gaussian filter (sigma = 1). 5. The image was auto-thresholded using the Li algorithm (Li and Tam, 1998)). 6. The mask was eroded three times to better align the mask boundary with the NMJ membrane.

#### Generation and evaluation of composite and individual PAZ segmentations

We considered three basic approaches to segmenting the PAZ. First, we considered segmenting individual PAZ channels into meshes. The advantage of this approach is that it allows for a comparison of architectures for individual proteins. However, this approach had limitations as well, since our goal was to conceptualize the PAZ as a unified membrane domain, in which many of these proteins physically interact at some point in time and space. Further, individual PAZ representations matched each other poorly, and made it difficult to directly compare different PAZ protein concentrations, colocalizations, and relationships to active zone properties. To solve these issues, we next considered segmenting the PAZ using the active zone as a fiduciary marker for the PAZ. This creates a unified PAZ representation which simplifies PAZ-PAZ and active zone-PAZ comparisons, but necessarily misses some features of the PAZ, such as PAZ units with multiple active zones. Finally, we developed a composite PAZ segmentation strategy that accounts for both the PAZ and active zone signals in each experiment (**Fig S1A**).

Segmentation of each NMJ into a PAZ mesh representation was performed as follows: 1) The image stack was background subtracted (rolling ball radius = 50 pixels). 2) Individual channels were normalized by the minimum and maximum intensities per channel within the masked NMJ regions. 3) The image was blurred using a Gaussian filter (sigma = 1). 4) (unique to the composite PAZ approach) PAZ channels were summed and the active zone channel subtracted to create a composite intensity image (resulting in a different image than the composite for NMJ segmentation because here we subtract the active zone signal). The composite image was then processed with an ‘Internal Gradients’ filter using an octagonal element of radius 5 (Legland et al., 2016). 5) Local minima were detected using an empirically optimized threshold value (see below). 6) Mesh regions were identified using a marker-controlled watershed algorithm (Legland et al., 2016). Individual channel segmentations were generated as above but using only single channels and with an internal gradient radius of For each experiment, the composite mesh was defined by the composite of two PAZ proteins and BRP. All pairwise PAZ groups were represented across all experiments, and we did not observe experiment to experiment variability in the segmentation or metrics (**Fig 3**), indicating that this strategy was robust to the specific PAZ channels used for segmentation. For analysis, the mesh region was considered a four pixel wide area around the PAZ unit boundary (six pixel wide for STED), and the core was considered to be the remaining area within the PAZ unit.

When evaluating the different segmentation strategies, we observed that individual PAZ meshes correlated poorly with one another (**Fig S1C-E**), reflecting the complex and heterogeneous distribution of endocytic proteins. The strictly active zone-based PAZ also correlated poorly with individual PAZ protein segmentation, and was over-segmented due to the presence of a significant number of meshes with multiple AZ. The composite mesh segmentation most consistently represented active zone and PAZ architecture (**Fig 3**). Thus, we chose to use a composite mesh segmentation for all analyses where we compare different PAZ proteins (**Fig 2-4**). See code for all details.

#### Segmentation of active zone mesh and objects

Segmentation of a mesh representation based on active zones (as discussed above) was performed as for single PAZ channel segmentations, except that local intensity maxima were detected rather than local intensity minima. Local maxima were used both to generate the active zone-based mesh and to localize and count active zones within PAZ meshes. For segmentation into active zones objects, these local maxima were used as seeds to generate objects by the seeded region growing ImageJ plugin (ImageJ-Plugins). Merged active zones were split by a distance transform watershed (Legland et al., 2016).

#### Optimization of threshold value for min/max detection

The primary flexible parameter used to tune the segmentation of PAZ mesh is the tolerance/threshold value for the detection of local minima. A higher tolerance leads to a coarser segmentation (larger and fewer meshes) while a lower tolerance leads to a finer segmentation (small and more numerous meshes). To ensure consistent segmentation from image to image and experiment to experiment, we devised an optimization protocol to determine this threshold in an objective unbiased fashion. For each independent experiment, composite meshes were generated for each image using a wide range of tolerance values. The mesh generated by each (excluding meshes at the extreme edges of the NMJ (see below)) was used to measure the mesh ratio and average mesh size. Each image was scored as the geometric mean of mesh ratio, mesh area, and a spatial penalty representing what fraction of the NMJ was effectively represented by the mesh. The spatial penalty was devised to prevent systematic over-segmentation of the NMJ. For each image, every tolerance value was ranked by score and the tolerance value with the best average rank was selected as the threshold for the entire experiment and applied to all images in that experiment. An identical process was used to optimize segmentation of individual channels.

#### Filtering of edge units

While generating and analyzing PAZ meshes, we observed a noticeable edge artifact in most images due to generating a flat 2D representation of a 3D ellipsoid. We observed this artifact as a higher intensity (due to the summation of fluorescence along the vertical NMJ membrane by the point spread function) and an apparent higher number of active zones (due to the projection of active zones along the vertical surfaces of the NMJ). To exclude these edge effects from our analyses, we excluded all mesh units that occupied the edges of the NMJ (which we defined as having an average distance map value of 300 nm from the NMJ edge).

## QUANTIFICATION OF PAZ ARCHITECTURE

Code used to perform all analyses in the manuscript is freely available https://github.com/rodallab/paz-analysis.

### Jaccard Index

Jaccard Index was calculated as the area of intersection of two meshes (with a mesh width of 2 pixels) divided by the area union of two meshes.

### Centroid Agreement

Centroid agreement quantifies the degree of overlap between two sets of points A and B (having number densities ρ^A^ and ρ^B^, respectively). First, we calculated the average distance δ^A,B^ from each point in A to its nearest neighbor in B, and the average distance δ^B,A^ from each point in B to its nearest neighbor in A. δ^total^ was calculated as the density-weighted average of δ^A,B^ and δ^B,A^, i.e. δ^total^ = (ρ^A^ δ^A,B^+ ρ^B^ δ^B,A^)/ (ρ^A^+ ρ^B^). As a negative control, we compared δ^total^ to the situation of independent sets of regularly spaced points. We calculated this quantity explicitly in the case of a null model in which A and B are each made up of points on a hexagonal lattice, with a random overall shift between the two lattices. We label the value obtained by this null model as δ^null^: it may be shown by direct calculation that a very good approximation is obtained by setting δ^null^ = 0.4 (ρ^A^/ √ (ρ^B^) + ρ^B^/ √ (ρ^A^))/(ρ^A^ + ρ^B^). We then define the centroid agreement as the relative difference from this null model: (δ^total^-δ^null^)/ δ^null^ (see **Fig 3D**).

### Radial Profiling

Radial profiles were calculated per mesh unit as the average of all intensity profiles drawn from each mesh pixel to the local intensity minimum of the mesh unit. To average across mesh units, distance values were normalized from 0 (mesh edge) to 1 (core minimum). To prevent artifacts due to small mesh units, we excluded from this analysis any mesh with an area less than 250 nm^2^.

### Haralick’s Entropy

Entropy was calculated using the ImageJ plugin ‘Texture Analyzer’ (Julio E. Cabrera). To calculate entropy, circular meshes were first transformed into rectangles using the “Straighten” function in ImageJ.

### Spottiness

We processed images with a commonly used spot detection filter (Laplacian of Gaussian), and calculated the magnitude of response to this filter as the root mean squared value of the filtered image (because the filter ranges from positive to negative across spot boundaries). Higher scores indicate a more punctate or spotty distribution of signal.

### Centroid Distributions

Centroid distributions were measured in Python using PySal and pointpats libraries. First an alpha shape for each image was calculated using the pooled local minima (PAZ channels or composite) and maxima (for active zone) per image. Ripley’s G was calculated for composite or active zone point distributions. To determine the nature of the observed distribution, we generated model random, regularly spaced, or clustered distributions, each with a density matching the observed density (ρ^obs^). Random and clustered were generated using pointpats from poisson and poisson cluster process, respectively. Clustered distributions were generated in two rounds, an initial ‘parent’ round with an initial density of (ρ^obs^)^1/2^. Next, a cluster radius *r* was defined as the area of the alpha shape divided by 512. Finally, each parent generated a sufficient number of ‘children’ within a random radius on the interval 0-*r* to match ρ^obs^. Regularly spaced distributions were generated by a Matern Type II process using code adapted from H. Paul Keeler https://hpaulkeeler.com/simulating-matern-hard-core-point-processes/. One caveat of this approach is that modeled distributions were generated using the alpha mask as the boundary, which can be slightly broader than the actual NMJ area.

### Colocalization

Pearson’s correlations were measured on 2D maximum intensity projection images. As observed in medial slices (Fig S1A), the majority of PAZ signal is localized to the plasma membrane within our (X,Y) imaging resolution. As our imaging modalities have even less resolution in the Z, the maximum intensity projection of our image half-stacks does not sacrifice any spatial information, while allowing us to specifically address the lateral organization of PAZ proteins and their colocalization in the plane of the membrane.

## STATISTICAL ANALYSIS

Graphs were prepared and statistical analyses performed using Python (using Matplotplib, Seaborn, Scipy-Stats, Scikit-Posthocs libraries). For normally distributed data, comparisons were made using either T-test or ANOVA with posthoc Bonferroni’s multiple comparisons test. For non-normally distributed data, comparisons were made using either Mann-Whitney U test, or Kruskal-Wallis test with posthoc Dunn’s test. All quantitative analyses presented were performed on datasets composed of two PAZ channels and BRP, imaged by Airyscan in SR mode. As we observed low between experiment variability (as shown on plots), statistical comparisons are made using pooled data from multiple experiments. For the specific proteins labeled in each individual experiment and the N of animals, neurons, and PAZ units analyzed in each independent experiment please see **Supplemental Table S1**. No specific power analyses were performed; sample sizes were chosen based on established protocols and statistical analyses for significance, as detailed for all experiments here. See ‘Supplemental File 1-Statistical Analyses.zip’ for each statistical test performed for each experiment presented in this study. Statistical significance denoted in all graphs * p<0.05, ** p<0.01, *** p<0.001.

## 4. Results

### Endocytic and adhesion proteins heterogeneously localize to the PAZ

As a model system, we used the *Drosophila* larval NMJ (**Fig 1A**), which consists of large (3-6 μm diameter) synaptic boutons containing multiple (10-30) active zones, allowing us to resolve the organization of PAZ molecules at synaptic membranes (see **Table 1** for a summary of genes and labeling strategies). We focused our analyses on synapses at rest, where the number of spontaneous exocytic and endocytic events is approximately five per bouton per minute (Akbergenova et al., 2018; Del Signore et al., 2021; Melom et al., 2013). We imaged PAZ protein distribution and architecture primarily by Airyscan microscopy, and confirmed results by STED microscopy (see **Fig 5** below). As previously observed in both live and fixed samples, we observed that PAZ proteins generally form a membrane-associated mesh surrounding active zones (**Fig 1B**, (Del Signore et al., 2021; Koh et al., 2004; Marie et al., 2004; Roos and Kelly, 1999; Sone et al., 2000; Winther et al., 2015)). We further noted in both live (**Fig 5** and Del Signore et al., 2021) and fixed preparations (**Fig S1A**) that the vast majority of signal localized to the plasma membrane (or a domain adjacent to the membrane within the resolution limits of our imaging), with little signal further from the membrane, where most synaptic vesicles are distributed (Denker et al., 2009). Thus, only a small fraction of PAZ proteins are likely sequestered in vesicle pools, though this has been a common model for their regulated deployment. We noted a surprising complexity to the distributions of individual PAZ molecules within the PAZ, with significant variation in local concentrations and a striking degree of non-colocalization of different pairs of proteins, even those that physically interact such as Dap160 and Nwk (Rodal 2008, O’Connor Giles 2008, Almeida-Souza 2018) (**Fig 1B-D**).

**Figure 1.**
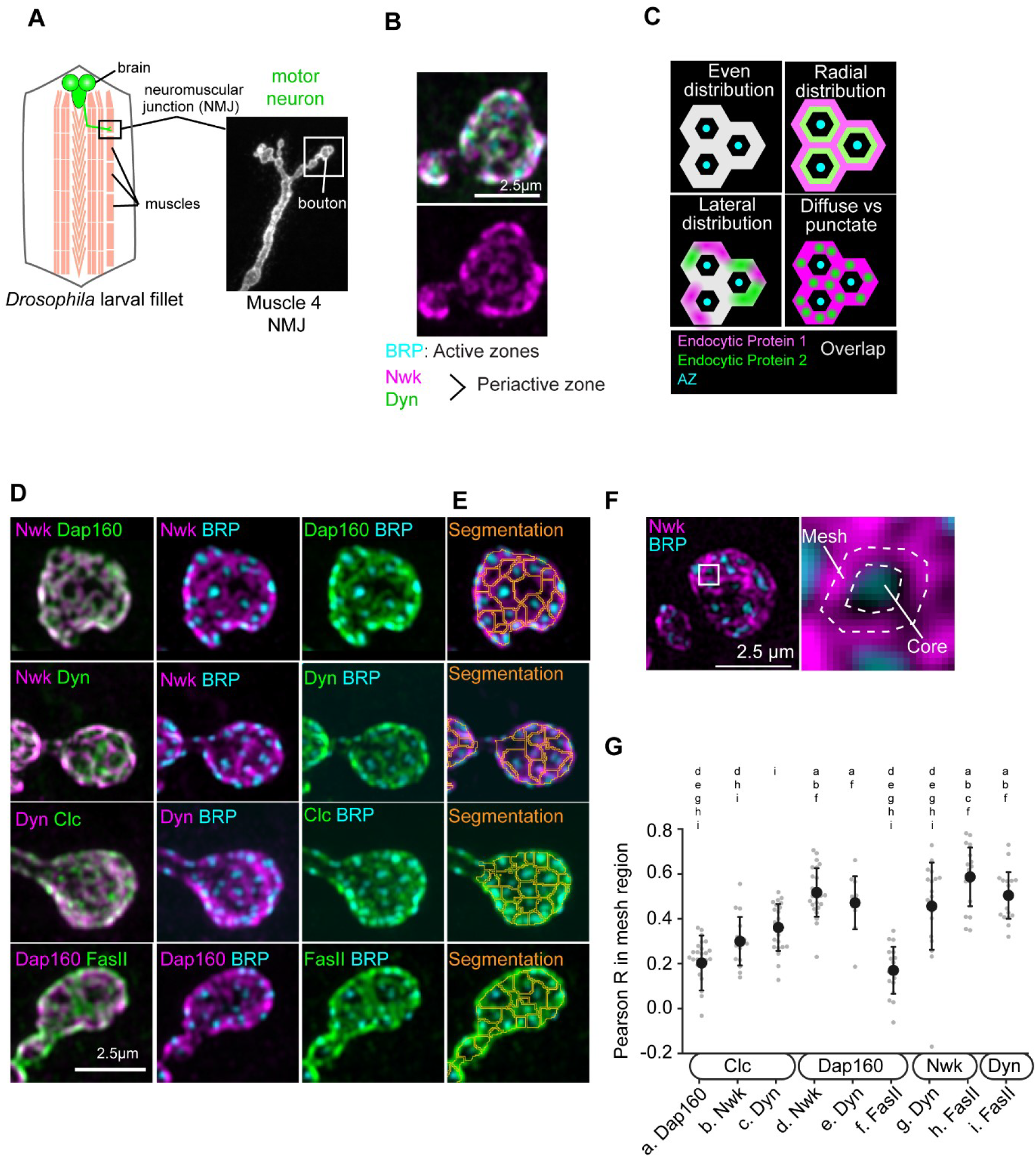
Endocytic proteins heterogeneously localize within the PAZ. (A) Schematic of the *Drosophila* NMJ prep. Left-cartoon illustration of the *Drosophila* larval fillet prep, with muscles highlighted in light red and the brain/motor neurons highlighted in green. Right-Example confocal image of a muscle 4 NMJ labeled with HRP. (B) Airyscan image of NMJ labeled with BRP (cyan), Nwk (magenta), and Dynamin (green). (C) Cartoon illustrating potential ways that endocytic proteins might differentially accumulate – radially (from core to mesh), laterally (within the plane of the mesh region), or relatively more diffuse or punctate. (D) Panel showing the range of patterns of accumulation of endocytic proteins at the NMJ, as labeled. Note that Nwk and FasII form a more highly restricted mesh, Dynamin frequently intercalates into the core region, and Clc accumulates in a more punctate pattern. (E) Example segmentations of the indicated channels into representative composite PAZ mesh. See Fig S1 and methods for more detail. (F) Schematic representation of a single PAZ mesh unit, divided into mesh and core regions. (G) Pearson R between the indicated pairwise comparisons within the mesh region indicates that colocalization of PAZ proteins is quite variable and ranges from low (∼0.2) to moderately high (∼0.6), but never approaches perfect or homogenous colocalization. Letters above each group indicate which groups are significantly different from the given column at p<0.05. Dots in G represent the average mesh value for a single image. Black dot and error bars indicate mean +/-Std Dev for all images. For this and subsequent figures, see **Supplemental Table S1** for a summary of N of animals, neurons, and PAZ units analyzed per experiment.

To quantify this complexity and gain insight into the architecture of the PAZ, we segmented the synapse into a set of PAZ units, where each individual PAZ unit represents a perimeter of PAZ proteins surrounding between 0-3 active zones (see Fig 3E below). For most experiments, we segmented PAZ architecture based on the combined fluorescence signal of two PAZ proteins and the active zone marker BRP (**Fig 1E, Fig S1A-F**; see methods for rationale and details). We then subdivided each PAZ unit into exocytic ‘core’ and endocytic ‘mesh’ regions (**Fig 1F**). For a subset of analyses (**Fig S1E-F, Fig 3C-D**) we also segmented PAZ units using individual PAZ or active zone signals. Lastly, we segmented each active zone into a discrete object to quantify its size and intensity. The respective sizes of these regions are on average: PAZ mesh .237 +/-.05 μm^2^; PAZ core .101 +/-.01 μm^2^; active zone objects .054 +/-.002 μm^2^ (see **Fig S1B** for distributions), indicating that the PAZ unit is roughly two thirds mesh and one third core, and that approximately half the core is occupied by the active zone marker BRP.

To formally test whether the PAZ is a heterogenous structure, we first quantified the colocalization of endocytic and adhesion molecules within the composite PAZ mesh (**Fig 1G**). If the PAZ is a homogenous structure, then we expect strong and consistent pairwise colocalization between PAZ proteins. Instead, we find that Pearson R values range from less than 0.2 to 0.6, indicating significant differences in where individual proteins accumulate within the PAZ. Strikingly, these weak and variable relationships were true even for pairs such as Dynamin and Dap160, two well characterized PAZ proteins that physically interact and whose levels, localization, and activity are coupled (Koh et al., 2004; Rodal et al., 2008; Roos and Kelly, 1998; Winther et al., 2013). As a negative control, we analyzed the colocalization of PAZ proteins with the active zone marker BRP in the mesh region (**Fig S1C**). Consistently for all PAZ proteins, the Pearson R to BRP was approximately zero, supporting the conclusion that even the lowest PAZ-PAZ colocalizations are above background. Together, these observations suggest a previously unappreciated complexity to the organization of the endocytic machinery at synaptic membranes.

### Proteins partition differently within the PAZ

We next asked precisely where and how PAZ proteins accumulate at individual PAZ mesh sites. We considered three ways that PAZ proteins might differentially localize – radially, by partitioning differentially between mesh and core regions; laterally, by accumulating in distinct subdomains within the PAZ mesh or core; and finally the extent to which their accumulation is diffuse versus punctate (see schematic **Fig 1C** and example images **Fig 2A**). To examine radial distribution, we generated a normalized radial profile extending from the mesh to the centroid of the core for each protein (see methods for details). As expected, we found that PAZ proteins were enriched in the mesh region, and gradually tapered in the core. Conversely, the active zone protein BRP was strongly enriched in the PAZ core (**Fig 2B**) and tapered in the mesh region. We noted that the degree of radial polarization was much lower for PAZ proteins compared to BRP, suggesting a non-trivial accumulation of PAZ proteins within the core region. Further, we noted small but significant differences in the degree of enrichment in the PAZ mesh between PAZ proteins-in particular, Dynamin had a significantly lower polarization than other PAZ proteins (**Fig 2B-C, Fig S2A**). To quantify these differences, we calculated the ‘mesh ratio’, which is the ratio of the mean mesh intensity over the mean core intensity, and found that both Nwk and FasII were significantly more polarized than Dynamin (**Fig 2D**). This is consistent with our qualitative observations that Dynamin frequently accumulated within the core region adjacent to the active zone (**Fig 2A**) and may reflect a wider range of roles for Dynamin in vesicle recycling, or a distinct mechanism of Dynamin recruitment relative to other endocytic proteins (see Discussion).

**Figure 2.**
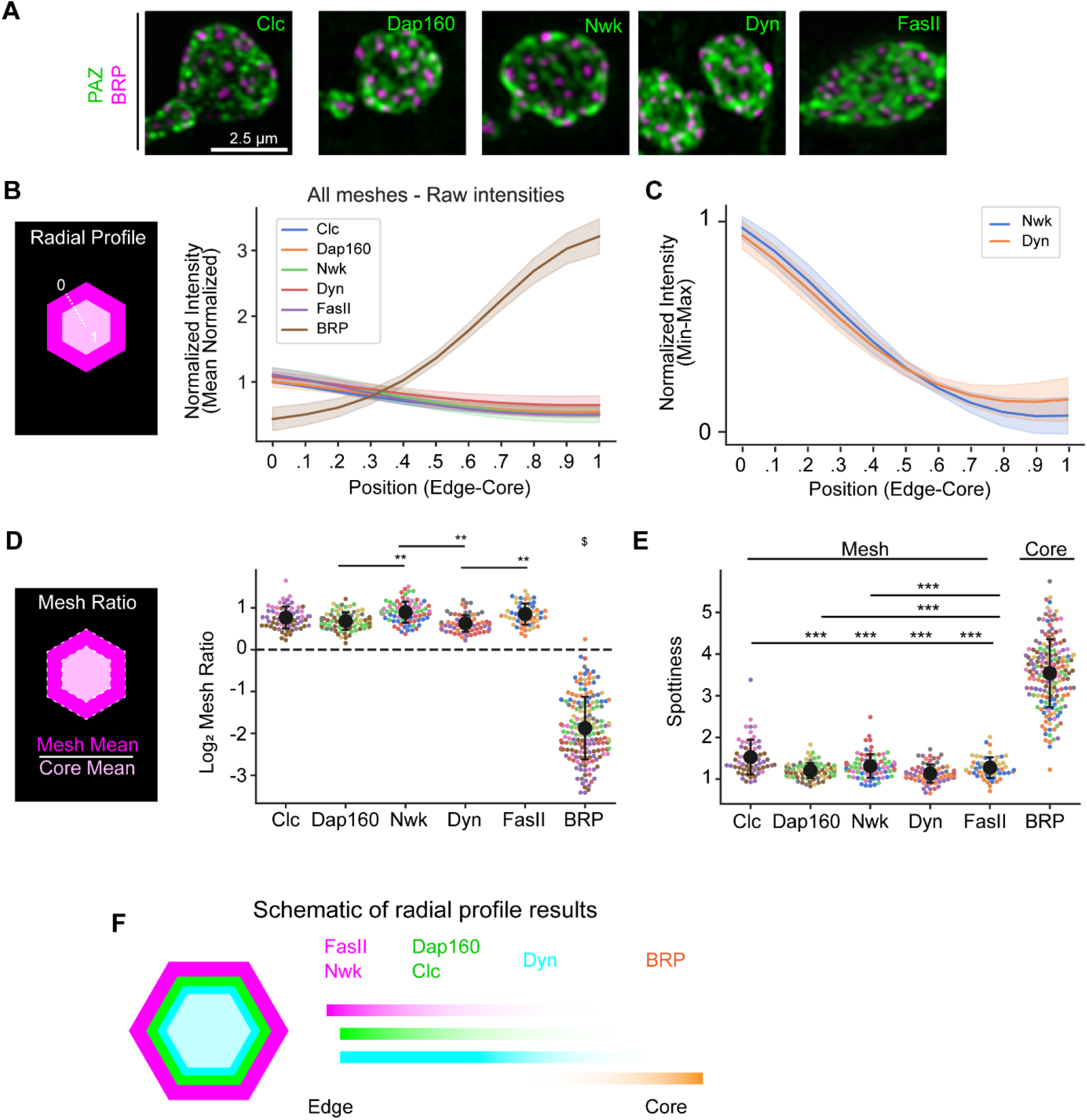
Endocytic proteins differentially distribute within the PAZ. (A) Example NMJ boutons labeled with BRP and single PAZ proteins to highlight differences in accumulation. (B-C) Radial profiles of BRP and PAZ proteins plotted against position from mesh edge (0) to core center (1), as depicted in the cartoon schematic. (B) Plots of mean normalized intensity by radial distance; BRP is inversely and more highly polarized compared to PAZ proteins. (C) Nwk and Dyn intensity (normalized min to max) plotted against radial position. Nwk is more highly polarized than Dyn, particularly at the core center. See Fig S2 for all other pairwise profiles. (D) Quantification of the mesh ratio: the mean intensity in the mesh region divided by the mean intensity in the core, as depicted in the cartoon schematic. Nwk and FasII are significantly more enriched in the mesh than Dyn. (E) Quantification of ‘spottiness’, measured as the root mean squared of each image processed with a Laplacian of Gaussian filter. Higher values correspond to more punctate signal. (F) Cartoon summarizing radial distributions of PAZ and active zone proteins. Lines in B-C indicate the mean +/-Std Dev mesh profile from 3-9 independent experiments per channel (see Supplemental Table 1 for details). $ in D indicates that BRP is significantly different than all other groups (p<.001). Dots in D-E represent the average mesh value for a single image; spot color indicates independent experimental replicates and does not correspond to colors in the legend in A. Black dot and error bars indicate mean +/-Std Dev for all images.

We next quantified how PAZ proteins accumulate within the mesh and core regions. We quantified the ‘spottiness’ of protein distribution using a commonly-used spot detection filter (see method for details, **Fig 2E**). Due to its purely punctate accumulation in the core region, BRP exhibited a much higher spottiness score than the PAZ proteins. Among PAZ proteins, we found that clathrin was significantly spottier than Dap160 and Dynamin (**Fig 2D**) within the mesh. We noted qualitatively that clathrin puncta were far more numerous than the expected number of endocytic events at this synapse. We also analyzed the pixel intensity distribution more generally, using Haralick’s Entropy (Haralick et al., 1973), **Fig S2B**), which quantifies signal heterogeneity over a local spatial window. Consistent with its condensed/punctate accumulation, Clathrin exhibited a significantly lower entropy score compared to the other PAZ proteins. Together, these data demonstrate that PAZ proteins exhibit both overlapping and distinct patterns of accumulation both radially and laterally at the PAZ (**Fig 2F**).

### PAZ architecture does not perfectly align with active zone distribution

A conventional assumption regarding the PAZ is that it is defined spatially by the placement and patterning of active zones at the synaptic membrane. We sought to test this by comparing the micron-scale architectures defined by BRP-labeled active zones and by our composite PAZ. We selected BRP as an active zone marker because it has been extensively characterized at this synapse in structural (Böhme et al., 2019; Fouquet et al., 2009; Kittel et al., 2006; Matkovic et al., 2013) and physiological studies (Akbergenova et al., 2018; Böhme et al., 2019; Melom et al., 2013), and at the resolution of our imaging forms a clear spot well suited for localization and intensity analyses. Functionally, BRP is a scaffold that is required for the maturation of active zones and the normal clustering of voltage gated calcium channels (Fouquet et al., 2009; Kittel et al., 2006). First, we analyzed the spatial distributions of the local maxima of BRP with the local minima of our composite PAZ signal. We quantified the spatial patterning using Ripley’s g function, which quantifies the fraction of points for which the nearest neighbor is within a given distance *d* (**Fig 3A-B**). The spatial patterning of BRP and composite PAZ spots were indistinguishable. To determine the nature of this distribution, for each image analyzed we also created model point distributions generated by either purely random (Poisson), clustered (Poisson clustered) or regularly spaced (Matern Type II) processes (**Fig 3A**; see methods for details and discussion of caveats). BRP and composite distributions most closely matched a regularly spaced distribution, with the minimum observed spacing between active zones of ∼325nm (**Fig 3B**). These data suggest that the relative placement of active zone/PAZ units is not random, and that mechanisms exist to regulate their spacing. Indeed these mechanisms may be linked, as PAZ mutants exhibit changes in active zone spacing (Dickman et al., 2006).

**Figure 3.**
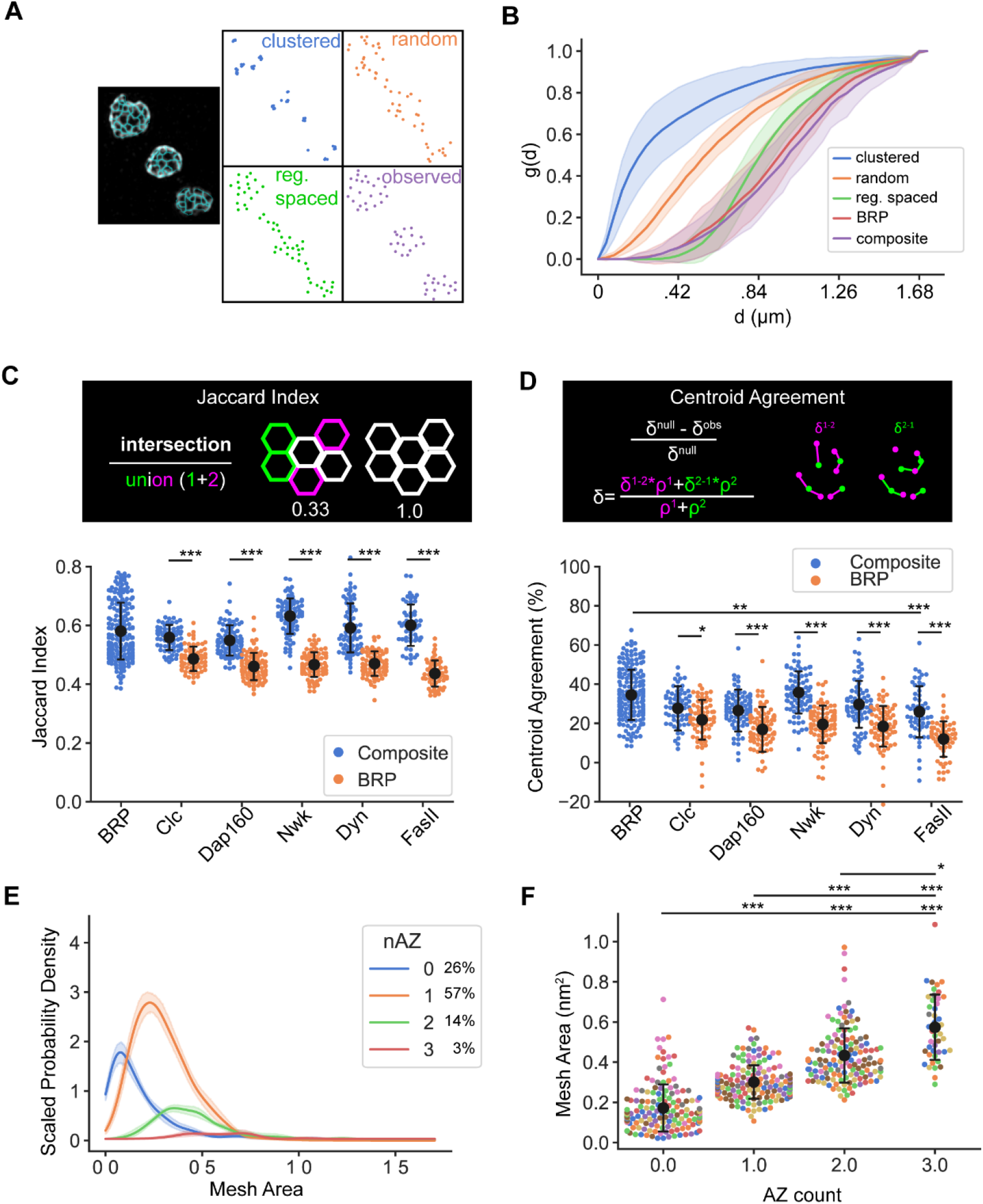
Active zone position and pattern only partially define PAZ architecture. (A-B) Example observed and model patterning of BRP and composite mesh centroids. (A) Left: Example NMJ with composite segmentation labeled in cyan. Right: Examples of modeled clustered, regularly spaced, and random spatial point distributions for the example NMJ, compared to the observed composite mesh centroid pattern. (B) Quantification of Ripley’s g function shows that spatial patterning of BRP and composite segmentations are indistinguishable, and best resemble a regularly spaced distribution. (C-D) Comparison of active zone and composite PAZ segmentations. Both Jaccard Index (C) and Centroid error (D) indicate that composite segmentations match active zone and individual PAZ meshes similarly well, and significantly better than active zone-based segmentations. (E) Probability density estimates of mesh areas from composite based segmentations, broken down by the number of active zones in the mesh. Active zone number correlates with mesh size. (F) Quantification of mesh area of composite meshes containing 0-3 active zones. The area of meshes significantly increases as active zone number increases. Dots in C,D represent the average value for a single image, black dot and error bars indicate mean +/-Std Dev for all images. Lines in B, E represent the mean +/-Std Dev per image (B) or experiment (E). Dots in F represent the average value for a single image, colors represent measurements made from independent experiments. black dot and error bars indicate mean +/-Std Dev for all images.

We next asked to what extent the actual architectures defined by BRP versus composite meshes agree with each other, and with the meshes generated by segmentation of individual PAZ channels. We analyzed architecture agreement by Jaccard Index (**Fig 3C, Fig S3A**,**C**) as well as a custom metric called centroid agreement, which measures the overlap between two sets of points by analyzing the average nearest neighbor distance between each set of points, compared to a theoretical null value for two sets of random points (**Fig 3D, Fig S3B**,**D**). Both measurements indicate that BRP and composite architectures agree only moderately. Further, we note that the composite mesh matches BRP and individual PAZ meshes similarly well, while the BRP mesh is significantly worse at representing each individual PAZ signal (compare blue to orange dots in Fig **3C-D**). Together, these data indicate that PAZ architecture reflects both active zone-dependent and independent features, and that the PAZ architecture is not purely defined by the active zone.

An obvious and intriguing example of the differences between these architectures is the existence of composite PAZ units containing zero or multiple active zones (**Fig 3E-F**). We noted that while the majority of PAZ meshes contained exactly one active zone (57%), many contained zero (26%), and a small fraction contained multiple (17%). Further, we noted that PAZ mesh size scaled with active zone number (**Fig 3F**). The assembly and composition of active zones changes developmentally (Akbergenova et al., 2018) and homeostatically (Böhme et al., 2019; Hong et al., 2020; Weyhersmüller et al., 2011), including the formation of multiple linked active zones (Hong et al., 2020). It will be interesting to determine the physiological properties of these multi-active zone PAZ units, and to what extent these represent transient developmental or stable architectural features.

### PAZ architecture correlates with active zone properties

We next asked whether PAZ domain architecture correlates with the properties of nearby active zones. Active zones at the *Drosophila* NMJ can vary >50-fold in their probability of vesicle release (Akbergenova et al., 2018; Melom et al., 2013). We noted that PAZ proteins also exhibited significant variability between PAZ mesh units, with a standard deviation of ∼25% of the mean intensity of each protein (**Fig S4A**). Here we asked whether the intensities and distributions of PAZ proteins covaried with active zone intensity (see schematic **Fig 4A**). This analysis takes advantage of the fact that the concentration of the scaffold protein BRP positively correlates with calcium channel numbers and active zone release probability (Akbergenova et al., 2018; Gratz et al., 2019; Melom et al., 2013). To simplify the analysis, we focused on PAZ meshes containing exactly one active zone. We compared BRP intensity to 18 PAZ measurements including PAZ area, PAZ protein concentration and distribution, and protein colocalization. Many parameters exhibited a significant correlation to active zone intensity (**Fig 4B-E, Supplemental Table 2**). Most notably, we found that the concentration of PAZ proteins (measured as their mean intensity) in both core and mesh regions strongly correlated with active zone intensity, suggesting that the concentration of PAZ proteins increases as release probability increases. Notably, for the lone adhesion protein in our analysis (FasII), this correlation was either not observed (in the core region, **Fig 4C**) or strikingly lower (in the mesh region, **Fig 4D**), suggesting this scaling may either be a specific property of endocytic PAZ proteins or perhaps reflects different levels of dynamics between the membrane associated endocytic proteins and the transmembrane FasII. We also noted that the mesh ratio of Clc, Dap160, and Nwk exhibited a significant negative correlation with active zone intensity (**Fig 4B, Fig 4E**), while Dynamin and FasII do not, indicating that only a subset of endocytic proteins redistribute according to the composition or physiology of nearby active zones.

**Figure 4.**
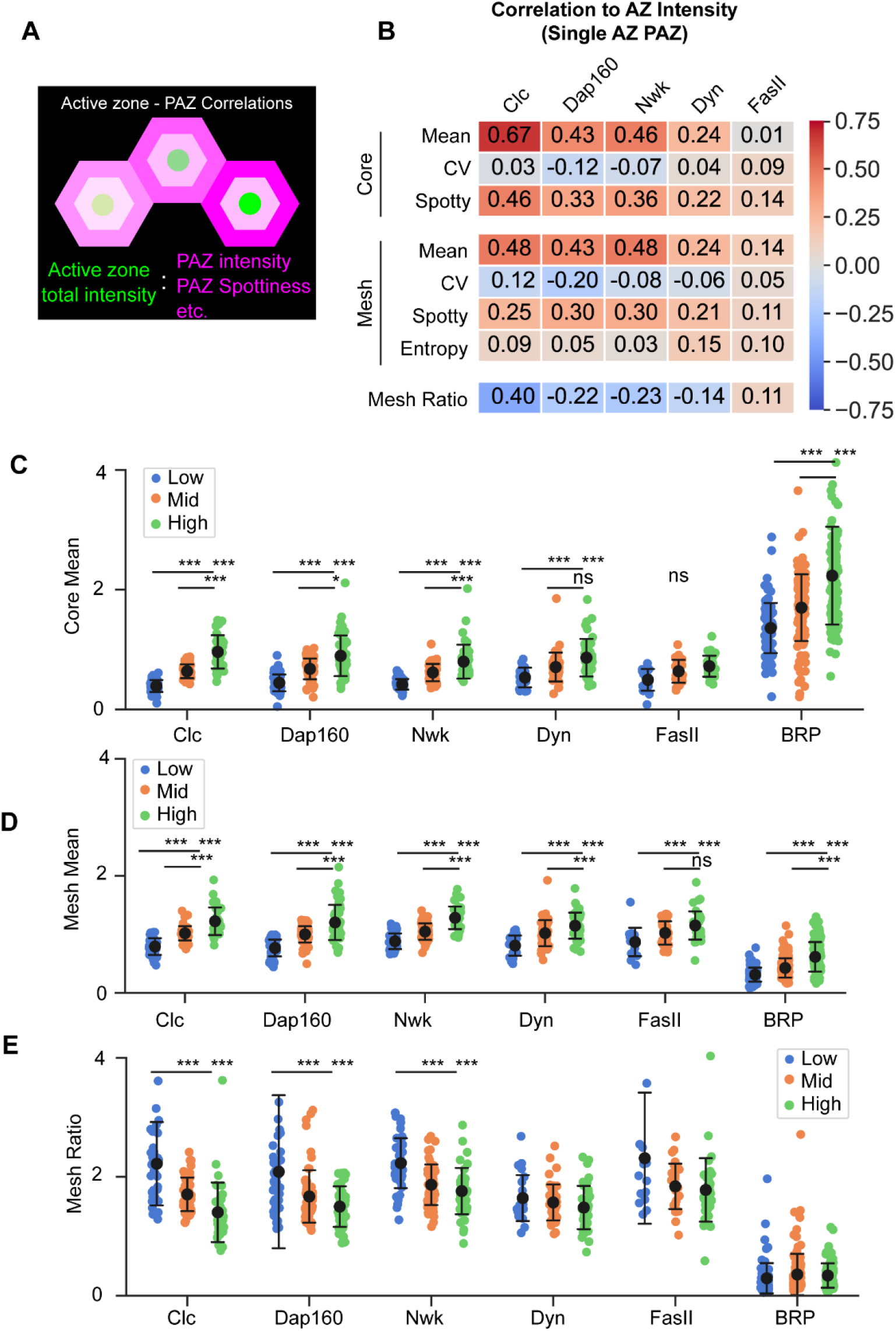
Synaptic endocytic protein architecture correlates with active zone structural properties. (A) Schematic of comparisons between active zone intensity and PAZ protein distributions. (B) Table summarizing the Pearson correlation between active zone integrated density and the indicated PAZ metrics. Notably, accumulation of all PAZ proteins except FasII in core and mesh regions generally correlate with active zone intensity. Mesh ratio exhibits a variable but generally negative correlation to active zone intensity. Additional metrics shown in Table S2. (C-E) Comparison of core mean intensity (C), mesh mean intensity (D), and mesh ratio (E) for PAZ and active zone proteins binned by active zone intensity quantile. (C-D) Consistent with Pearson values, in most cases PAZ proteins exhibit significantly higher core means in the high active zone intensity bin, with FasII a notable exception. (E) Clc, Dap160, and Nwk show a clear decrease in mesh enrichment as active zone integrated density increases. Values in B represent the average Pearson R per experiment, calculated from individual meshes of all images. Dots in C-E represent the average value per image, color indicates the active zone integrated density bin. Black dot and error bars indicate mean +/-standard deviation of all images.

To validate our findings in fixed tissues, we measured the distribution and relationship of Nwk and BRP in living neurons. Nwk and BRP were labeled with HALO or GFP tags knocked in to the endogenous loci. We found that, as in fixed tissues, Nwk localized to a membrane proximal mesh surrounding active zones labeled with BRP (**Fig 5A-B**). Radial profiles and mesh ratios were quantitatively similar to those observed in fixed samples (**Fig 5C-D**), and relationships to BRP intensity were largely preserved (**Fig 5E-F**). We did note small differences in the relationship between Nwk CoV and Entropy with BRP intensity, which might reflect some degree of temporal integration during fixation. In total, however, we conclude that our results are robust to live versus fixed imaging.

**Figure 5.**
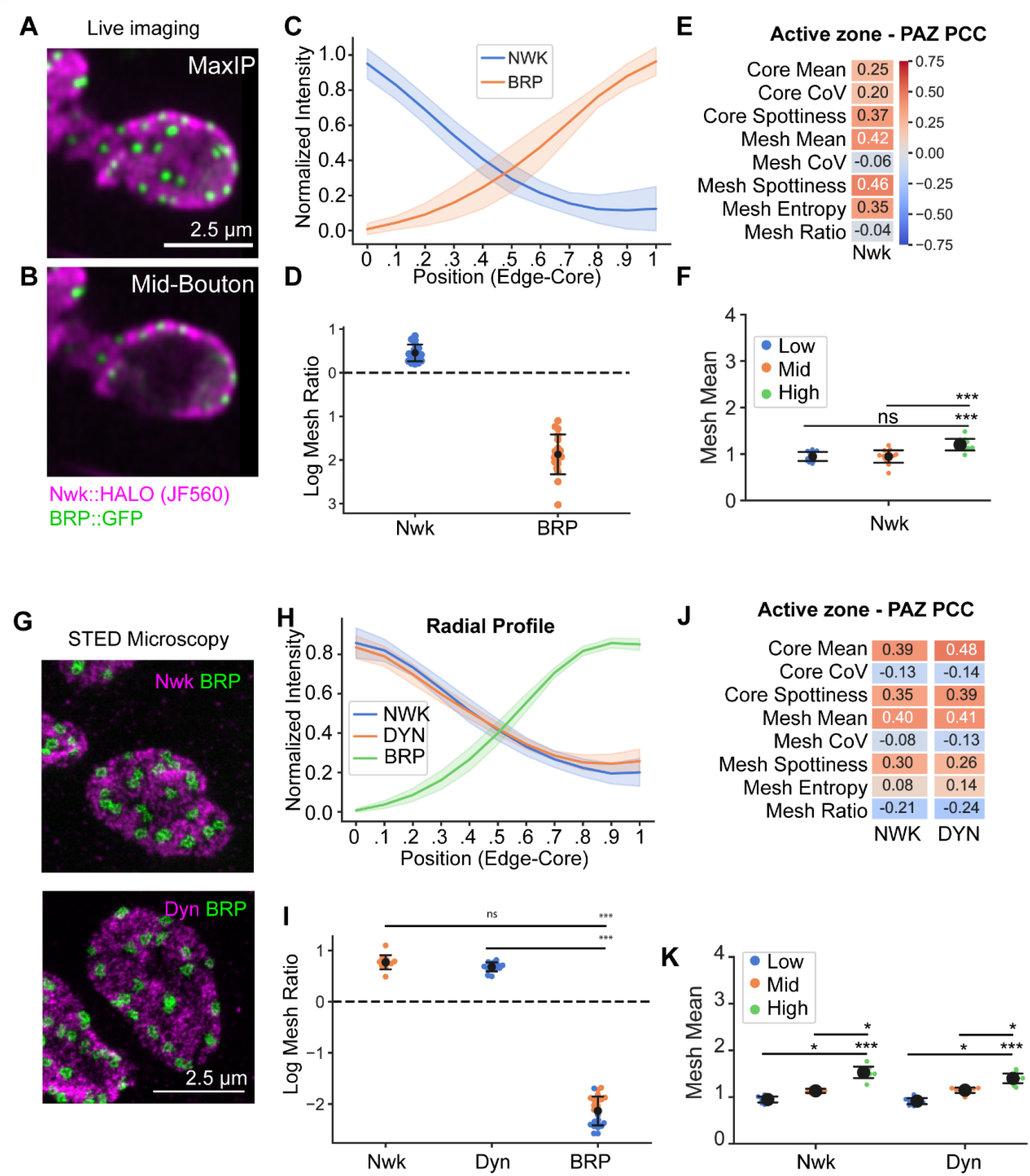
Live imaging and STED microscopy validate measurements of PAZ architecture. (A-B) Maximum intensity projection (A) and mid-bouton plane of a muscle 4 bouton labeled with endogenously tagged Nwk::HALO (JF549) and BRP::GFP. (C-D) Quantification of distribution of Nwk and BRP by radial profiling (C) and mesh ratio (D) show strong polarization of Nwk in the PAZ mesh and BRP in the PAZ core, consistent with observations in fixed tissues. (E-F) Comparison of Nwk and BRP intensity. Nwk accumulation and distribution correlate with BRP intensity, similar to results in fixed tissues. Correlations between BRP and Nwk CoV and Entropy differ slightly, suggesting features of the Nwk distribution may be sensitive to fixation. (G-K) Imaging and analysis of PAZ architecture by STED microscopy. Nwk-BRP and Dyn-BRP comparisons were made in independent experiments. (G) Single plane 2D STED microscopy images of Nwk and BRP (top) or Dyn and BRP (bottom). (H-I) Radial distribution by radial profile (H) or mesh ratio (I) demonstrates that Nwk and Dyn are enriched in the PAZ mesh, while BRP is strongly enriched in the PAZ core, consistent with Airyscan microscopy. (J-K) Quantification of active zone intensity – PAZ relationships. Similar to Airyscan, Nwk and Dyn accumulation in both mesh and core correlates with active zone intensity.

Finally, we tested the role of imaging modality on our analysis and conclusions. For these experiments we used STED microscopy to image Nwk or Dyn along with BRP (to compare active zone – PAZ distributions by STED, **Fig 5G-K**). In each case, we found that the observations made by Airyscan confocal microscopy were well supported by STED imaging (**Fig 5H-K**). STED microscopy better resolved differences in the distributions of the each of the proteins, such as the ring-like appearance of BRP labeled with the nc82 antibody. Notably, Dyn appeared as a dense collection of puncta (in contrast to the more continuous distribution by Airyscan), while Nwk appeared as a more continuous mesh, similar to that observed by Airyscan. In total, we conclude that our analysis pipeline can be readily applied to different super-resolution imaging modalities (and resolutions).

## 5. Discussion

### Pre-deployment model for synaptic endocytic machinery

Endocytic proteins accumulate at vastly higher concentrations and across much broader domains at synaptic membranes compared to non-synaptic membranes. Recent work suggests that a subset of endocytic proteins are pre-deployed to synaptic membranes (Del Signore et al., 2021; Imoto et al., 2021). Pre-deployment could provide an essential mechanism to allow for low latency vesicle recycling or release site clearance, to build a large pool of molecules for high-capacity cargo binding and internalization, or for other emergent requirements such as limiting cargo diffusion (Azarnia Tehran and Maritzen, 2022). This pre-deployment suggests that different regulatory mechanisms may control the molecular dynamics and activation of membrane remodeling at synapses compared to non-neuronal cells, and could result from multiple mechanisms including specialized lipid domains (Khuong et al., 2010; Li et al., 2020; Wahl et al., 2016) or protein phase separation (Imoto et al., 2021; Kozak and Kaksonen, 2022). Yet to date most studies of synaptic endocytic protein dynamics have focused on global transitions between membrane compartments such as the synaptic vesicle pool and the plasma membrane (Bai et al., 2010; Denker et al., 2011; Li et al., 2020; Winther et al., 2015, 2013) or general diffusion (Reshetniak et al., 2020; Wilhelm et al., 2014), and there remains no data that show how the spatio-temporal organization of endocytic proteins might drive synaptic vesicle recycling. Our work provides the first quantitative analysis of the micron-scale architecture and nano-scale distribution of a subset of the endocytic machinery at a synapse, and will be of great utility for understanding the distributions and regulation of dozens of additional components in future mechanistic studies. These analyses greatly extend prior qualitative descriptions of the micron-scale PAZ in both flies (Coyle et al., 2004; González-Gaitán and Jäckle, 1997; Marie et al., 2004; Rodal et al., 2008; Sone et al., 2000) and cultured mammalian neurons (Gerth et al., 2017; Wilhelm et al., 2014). Using our new method, we identify a novel and unexpected heterogeneity in the distribution of endocytic regulators *within* the PAZ and also between different active zones.

### Spatial segregation of PAZ components

The dense accumulation of PAZ proteins renders it difficult to assign discrete molecular activities or synaptic functions to specific membrane domains or protein accumulations. Our approach addresses this issue by providing a granular, spatially-resolved map of PAZ protein accumulation. We found that endocytic proteins exhibited distinct patterns of accumulation within the PAZ. Endocytic proteins differentially partitioned between active zone core and PAZ mesh domains (**Fig 2D**), localized with only partial overlap within the PAZ mesh (**Fig 1G**), and accumulated into relatively more diffuse or punctate patterns (**Fig 2E, Fig S2B**). It will be interesting to determine the molecular composition and function of these distinct patterns: does punctate clathrin reflect pre-deployed but immature clathrin structures or assemblies (Brach et al., 2014; Henne et al., 2010; Wood and Smith, 2021)? Does stronger Dyn accumulation in the PAZ core reflect its roles in modes of endocytosis that occur proximal to release sites, including ultrafast endocytosis (Kidokoro, 2006; Koenig and Ikeda, 1996; Kosaka and Ikeda, 1983; Kuromi et al., 2010; Teng and Wilkinson, 2000; Watanabe et al., 2013a)? Are more peripheral PAZ proteins held in a ‘poised’ or inactive state, or are they engaged in more diverse synaptic functions beyond synaptic vesicle endocytosis (Blanchette et al., 2022; Gimber et al., 2015; Jäpel et al., 2020; O’Connor-Giles et al., 2008; Rodal et al., 2011, 2008)? Alternatively, this heterogeneity could also reflect different temporal stages of a single biological process (such as stages of endocytosis). Finally, it will be important to investigate how spatial cues that regulate endocytic organization and function (such as lipid composition) are patterned in the PAZ (Khuong et al., 2010; Li et al., 2020; Wahl et al., 2016). Our new quantitative framework represents a first essential step to describe the distributions of these proteins at synapses; future experimental studies that employ these strategies will help us understand their regulation and function at synapses. For example, our method will be useful for identifying where and how endocytic proteins dynamically localize to active endocytic membrane remodeling events, e.g. by including markers of actin filament polymerization as a proxy for membrane invagination (Del Signore et al., 2021).

### Correlation of active zone and PAZ architecture

We found that the architecture and distribution of the endocytic machinery correlates with the functional properties of nearby sites of exocytosis (**Fig 4B-E**). It remains to be determined whether the coordination between PAZ and active zones in our experiments reflects acute or developmental adaptation of the endocytic machinery to vesicle release, or conversely, a role for the endocytic machinery in the assembly of active zones. Either may be likely as there are multiple examples of bidirectional cross-talk between active zones and PAZ: First, calcium influx via active zone-associated channels is thought to regulate endocytosis (Jorgensen et al., 1995; Littleton et al., 2001; Nicholson-Tomishima and Ryan, 2004; Poskanzer et al., 2006, 2003; Wu et al., 2009; Yao et al., 2011). Second, membrane lipids may coordinately regulate synaptic vesicle exocytosis (Honigmann et al., 2013; Khuong et al., 2013; Maritzen and Haucke, 2018; Walter et al., 2017) and endocytosis (Cremona et al., 1999; Di Paolo et al., 2004; Posor et al., 2013; Verstreken et al., 2003). Third, endocytic mutants increase the density of active zones at synaptic membranes (Dickman et al., 2006; Goel et al., 2019), supporting an instructive role for the PAZ machinery in active zone development. Finally, endocytic proteins play moonlighting roles in rapid clearance of cis-SNARE complexes from release sites (Hosoi et al., 2009; Hua et al., 2013; Jäpel et al., 2020; Kawasaki et al., 2000; Neher, 2010; Sakaba et al., 2013; Wu et al., 2014) to allow for high frequency exocytosis. Our analytical framework provides new opportunities to directly quantify the spatial and temporal relationship between exocytosis and endocytic molecular dynamics both during development, and in mutants that perturb synaptic activity and endocytic function.

## 6. Author Contributions and Notes

Conceptualization SJD, TGF, AAR Methodology SJD, TGF

Software SJD, TGF

Formal Analysis SJD, TGF

Investigation SJD, MGM, AMS

Data curation SJD, AAR

Writing SJD, TGF, AAR

Original Draft Writing SJD, TGF, AAR

Review & Editing SJD, MGM, AMS, TGF, AAR

Visualization SJD

Supervision AAR

Project Administration AAR

Funding Acquisition TGF, AAR

The authors declare no conflict of interest.

This article contains supporting information online.

## 7. Acknowledgments

The authors would like to thank Karsten Bahlmann and Mary Grace Velasco for assistance with STED imaging, Pascal Kaeser and Javier Emperador-Melero for helpful feedback on the manuscript, Hannah Yevick for helpful discussions, Graeme Davis for antibodies, the Bloomington *Drosophila* Stock Center (Indiana University, Bloomington, IN, NIH P40OD018537) for providing fly stocks, and the DSHB (created by the NICHD of the NIH) for monoclonal antibodies. This work was supported the Brandeis NSF MRSEC Bioinspired Soft Materials (NSF-DMR 2011846) and by R01 NS116375 (AAR and TGF).

## 8. Supplemental material

Supplemental material contains supplemental figures 1-4 and supplemental tables 1-2.

