## Supplemental material for "Quantitative mapping of synaptic periactive zone architecture and organization"

Figure S1

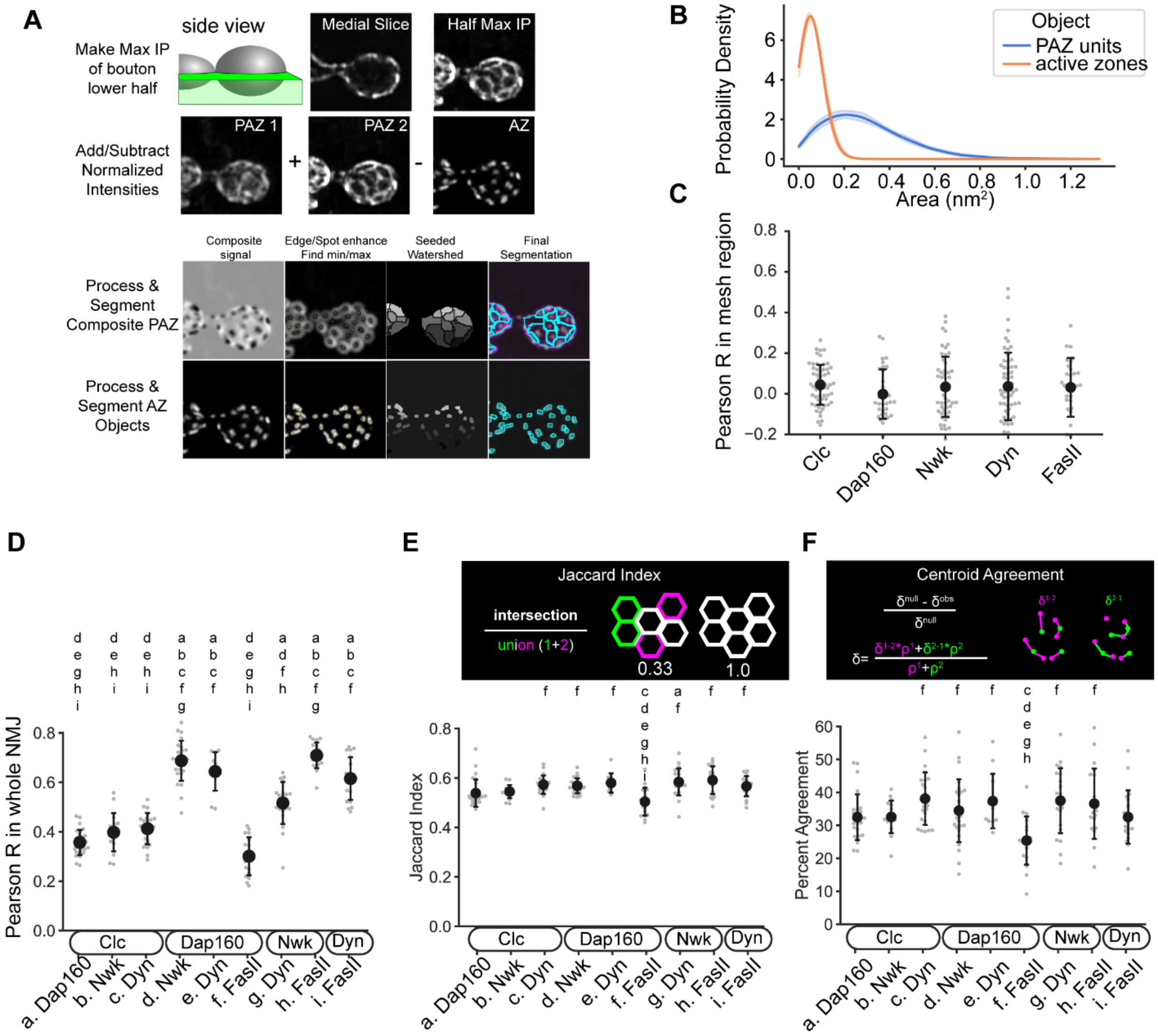


**Supplemental Figure *1.* Segmentation of PAZ architecture**. (A) Visualization of PAZ segmentation workflow. First, we generate maximum intensity projections of boutons from the medial cross section to the ventral surface of the neuron (i.e. the surface that is more deeply embedded in the muscle). To generate a composite segmentation, we normalize all channel intensities and add (PAZ) and subtract (active zone) channels to generate a single grayscale intensity map representing the NMJ. This image is then processed to enhance edge detection and images are segmented by a watershed algorithm using local minima as seeds. (For active zone object segmentation, local maxima are used as seeds followed by an additional distance transform watershed to separate touching objects.) (B) Kernel density estimates of the size distribution of composite PAZ units (blue) and active zone objects (orange). (C) Pearson R in the PAZ mesh region between BRP and individual PAZ proteins. The correlation is consistently near zero. (D-F) Quantification of PAZ architecture similarity by Pearson R (whole NMJ), Jaccard Index (D) or centroid nearest neighbor agreement (E) of pairwise PAZ protein distributions (D) or segmentations (E-F) within the same experiments. All measures suggest that PAZ proteins accumulate in partially distinct patterns across the synapse. Letters above each group in D-F indicate any groups significantly different at $p<.05$. Dots in D-F represent one NMJ or the average mesh value for a single NMJ. Lines show mean +/- standard deviation of all images.

Figure S2

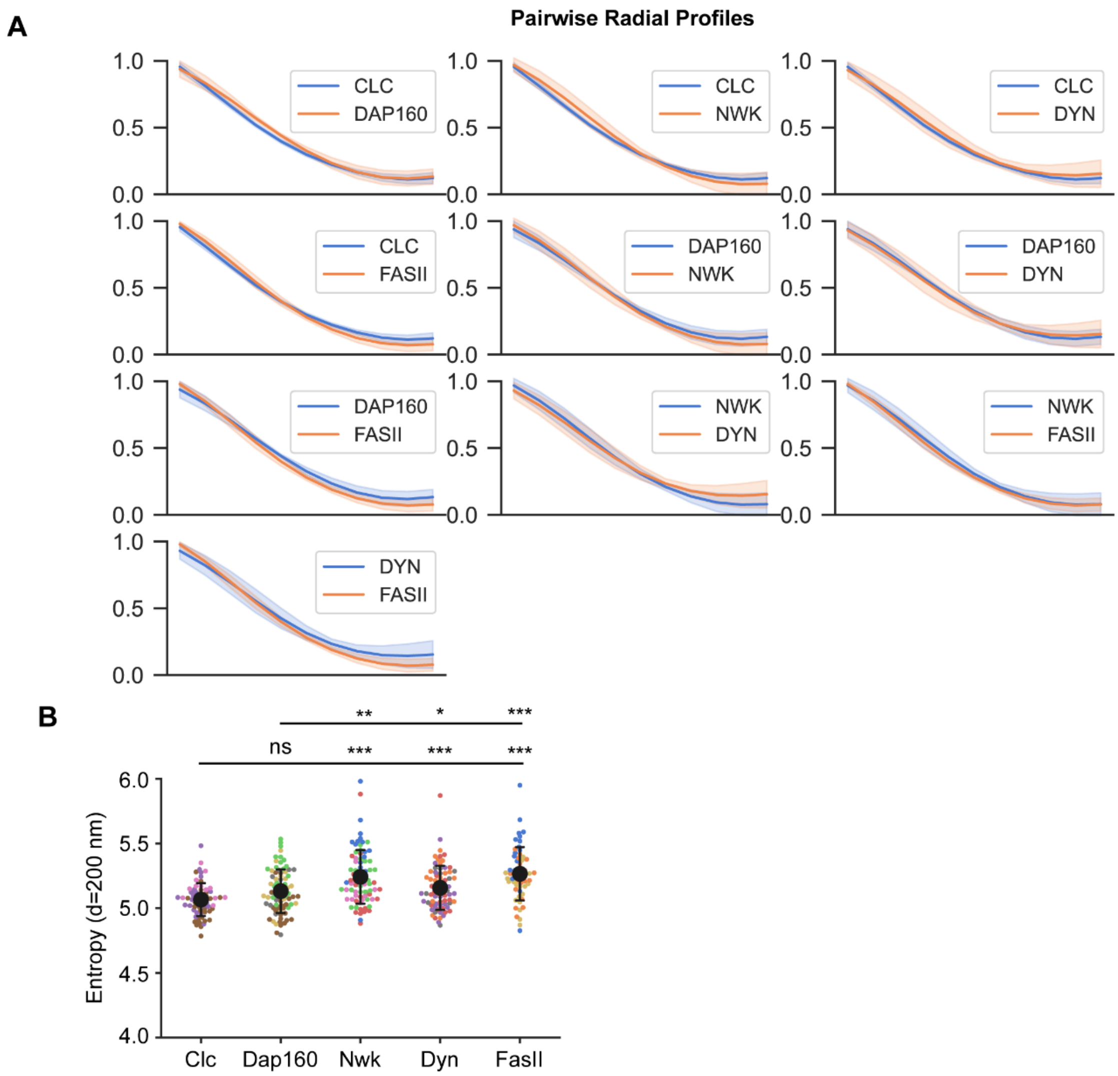


**Supplemental Figure 2**. (A) Pairwise radial profiles comparing PAZ protein distributions, same data as Fig 2A. Lines indicate the mean +/- Std Dev mesh profile from 3-4 independent experiments per channel (see Supplemental Table 1 for details). (B) Quantification of Haralick's Entropy in PAZ mesh regions, at a distance of 4 pixels. Clc exhibits a significantly lower entropy than Nwk, Dyn, or FasII.

Figure S3

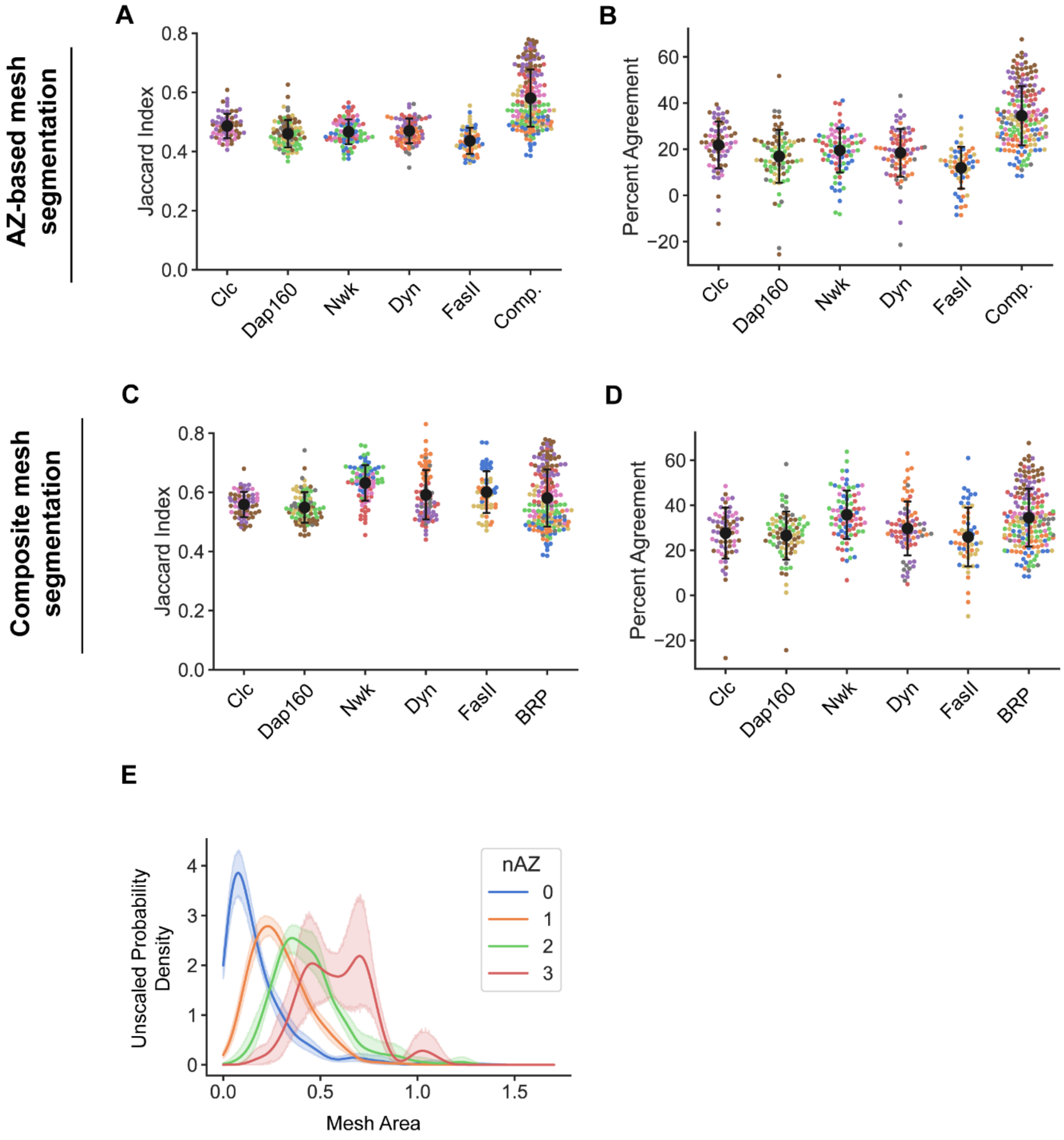


**Supplemental Figure 3. Active zone position and pattern partially define PAZ architecture**. (A-D) Alternative visualizations of the data shown in Fig 2 C-D comparing active zone and composite segmentations of the PAZ. Dots represent the average value for a single image, colors indicate independent experimental replicates, and black dot and error bars indicate mean +/- Std Dev for all images. (E) Unscaled probability density plot of the data shown in Fig 2F.

Figure S4

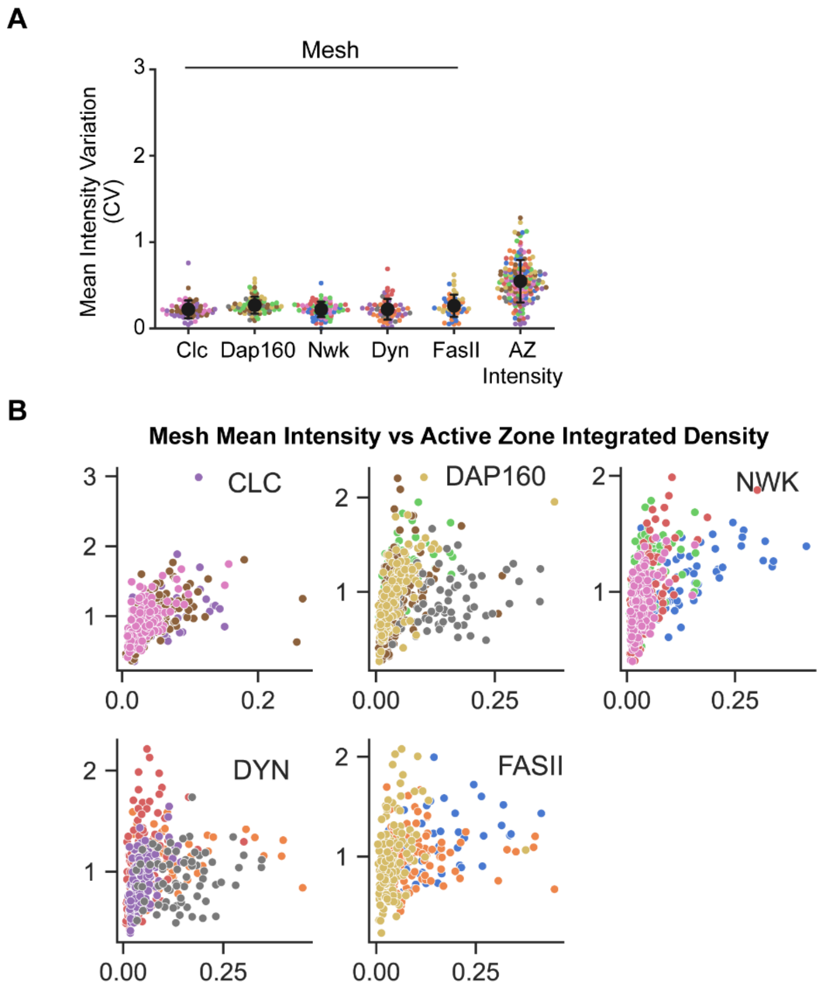


**Supplemental Figure 4.** (A) Quantification of intensity variation (Coefficient of Variation, CV) among mesh units. PAZ proteins (quantified in the mesh region) show consistent levels of between-mesh variation, while active zone integrated density exhibits significantly greater variation. Dots represent the average mesh value for a single image; spot color indicates independent experimental replicates. Black dot and error bars indicate mean +/- Std Dev for all images. (B) Scatterplots of mesh mean intensity plotted against active zone integrated density. Each spot represents one mesh, and differently colored spots are drawn from independent experiments.

| C1 | C2 | C3 | N Larvae | n NMJs | n PAZ units | n edge-filtered |
|---|---|---|---|---|---|---|
| NWK (970) | FASII (1D4) | BRP (Mimic) | 8 | 19 | 536 | 187 |
| DYN (Estes) | FASII (1D4) | BRP (Mimic) | 8 | 19 | 535 | 193 |
| Dap160 TSTEP | NWK (970) | BRP (nc82) | 5 | 26 | 1076 | 407 |
| Nwk (Mimic) | DYN (BD #41) | BRP (nc82) | 5 | 21 | 777 | 255 |
| DYN (Estes) | Clc (GFP) | BRP (nc82) | 6 | 23 | 616 | 181 |
| DAP160 (Roos) | Clc (GFP) | BRP (nc82) | 6 | 26 | 1176 | 374 |
| NWK (970) | Clc (GFP) | BRP (nc82) | 5 | 16 | 756 | 249 |
| DAP160 (Roos) | DYN (BD #41) | BRP (Mimic) | 6 | 11 | 435 | 144 |
| DAP160 (Roos) | FASII (1D4) | BRP (Mimic) | 6 | 18 | 823 | 273 |
| NWK (970) | BRP (nc82) | -- | 3 | 11 | 1364 | 968 |
| DYN (Estes) | BRP (nc82) | -- | 3 | 15 | 1689 | 1199 |

**Supplemental Table 1**. Summary of experimental replicates and labeling strategies.

**Supplemental Table 2**. Summary of all correlations measured between PAZ properties and active zone integrated density. A subset of these is shown in graphs in **Figure 4 B-E**.
